# Network integration shapes the loss and recovery of dynamical repertoires in disorders of consciousness

**DOI:** 10.64898/2026.06.26.734644

**Authors:** Fernando Lehue, Iván Mindlin, Carlos Coronel-Oliveros, Enzo Tagliazucchi, Jacobo D. Sitt, Patricio Orio

## Abstract

Disorders of consciousness (DoC) are associated with large-scale alterations in brain dynamics, yet which structural features constrain these changes remains unclear. We characterized dynamic functional connectivity (dFC) in 13 healthy controls (CNT), 11 minimally conscious (MCS) and 10 unresponsive (UWS) patients using two observables: the distribution of jump lengths between consecutive dFC frames and the occupancy of recurrent connectivity states. Controls showed more variable and temporally persistent jump lengths, captured by a lower Weibull shape parameter, and occupied states less constrained by structural connectivity. We then fitted a whole-brain dynamic mean-field model to each group and perturbed it node by node. Removing a core of ten posterior medial nodes with the highest integration (closely related to eigenvector centrality) moved the healthy-fitted model toward patient-like dynamics, whereas increasing the excitability of the same nodes moved the UWS-fitted model toward healthy-like dynamics. Other nodes had much smaller effects. These results extend previous lesion studies to bidirectional perturbations anchored to patient data, and suggest that a posterior core constrains the dynamical repertoire lost in DoC.

## Introduction

Disorders of consciousness (DoC) present a significant challenge in clinical neuroscience, characterized by profound alterations in awareness and responsiveness following severe brain injury [1]. DoCs are commonly classified using the Coma Recovery Scale [2] in the Minimally Conscious State (MCS) when there are traces of volitional responses to external stimuli, or Unresponsive Wakefulness Syndrome (UWS, also known as the Vegetative State) when there are none [3].

A key approach to understanding these alterations has been the study of large-scale patterns of spontaneous brain activity, which reflect the coordinated dynamics of distributed neural systems even in the absence of external stimulation. In DoC patients, such patterns are systematically disrupted, offering a neurophysiological correlate of impaired consciousness [4]. Functional Connectivity (FC) is defined as the statistical correlations between brain areas, usually when no explicit task is being performed (resting state). Although reduced activity in specific cortical regions (e.g., the precuneus and cingulate cortex) was long recognized as a hallmark of UWS [5], later imaging work showed that UWS is strongly associated with disruptions in FC in large scale networks of the resting brain [6] [7], especially the Default Mode Network (DMN). The DMN is a group of brain areas whose FC is intricately linked with processes such as self-referential thinking, episodic memory recall, goal-directed cognition, self-projection, and that has been central in studies of consciousness [8] [9].

Nowadays, it is accepted that consciousness does not rely on a single cortical area or network but requires brain-wide communication [10], and whole-brain FC alterations in DoC have provided insights and correlates of DoC’s etiology and mechanisms [11] [12]. In this regard, Demertzi et al. [4] demonstrated that healthy individuals (CNT) and DoC patients can be distinguished by systematic changes in the occupancy of dynamic patterns of FC.

Although static FC and dynamic FC (dFC) have been used to characterize the role of brain areas and their relation to consciousness [13] [14], the Structural Connectivity (SC) features that are crucial for explaining the role of each brain area on dynamic brain states are still not fully understood. Advancements in computational neuroscience and graph theory offer powerful tools to model brain dynamics and investigate the impact of SC features on FC [15] [16].

Whole-brain computational models offer a principled framework to go beyond characterization and probe causal mechanisms. By fitting models of this type to empirical FC or dFC, previous work has shown that DoC patients require lower global coupling to reproduce their spontaneous dynamics [17, 18], and that virtual lesions can recapitulate features of pathological brain states [19]. In silico stimulation can also force transitions between brain states [14], including out of DoC-like regimes [20]. Which nodes matter most is largely predictable from structure: in models of the healthy connectome, removing hubs and posterior midline regions has the largest and most widespread effects on simulated dynamics [21] [22], and out of nine graph-theoretical node measures, only eigenvector centrality and connector-hub status predicted how a lesion changes global synchrony and metastability [23]. However, these structural principles were established in models of healthy dynamics that were not fitted to patient data. It remains unclear whether they explain the dynamical changes measured in DoC patients, and whether the same structurally central nodes that push a healthy-fitted model toward patient-like dynamics when removed can also restore healthy-like dynamics when stimulated.

Here we ask how the structural connectivity profile of individual brain areas shapes their causal influence on whole-brain dynamics, using virtual lesions and local perturbations in a computational model fitted to empirical dFC measures in CNT and DoC patients. To this end, we use a whole-brain computational model that simulates virtual brain lesions and quantifies their impact on dFC. Specifically, we link a graph-theoretical integration measure, computed from the SC matrix, to the causal influence each node exerts on brain-wide dFC when it is disconnected or locally stimulated. This approach contributes to a mechanistic understanding of the structural underpinnings of dFC and their relevance to disorders of consciousness.

We first characterize the empirical dFC of CNT, MCS and UWS individuals with two observables (see Fig 1): the distribution of jump lengths between consecutive dFC frames, which describes how fast and how variably FC reconfigures, and, following [4], the occupancy of recurrent connectivity states obtained with K-means clustering, which shows that patients spend more time in states that resemble the SC. We then fit a whole-brain dynamic mean-field model to both observables in each group and perturb it node by node, removing each node from the CNT-fitted model and increasing the excitability of each node in the UWS-fitted model by reducing its local feedback inhibition. Finally, we relate the effect of each perturbation to the node’s integration, computed from the SC matrix with Hierarchical Modular Analysis [24, 25], and compare it with other centrality measures.

**Fig 1.**
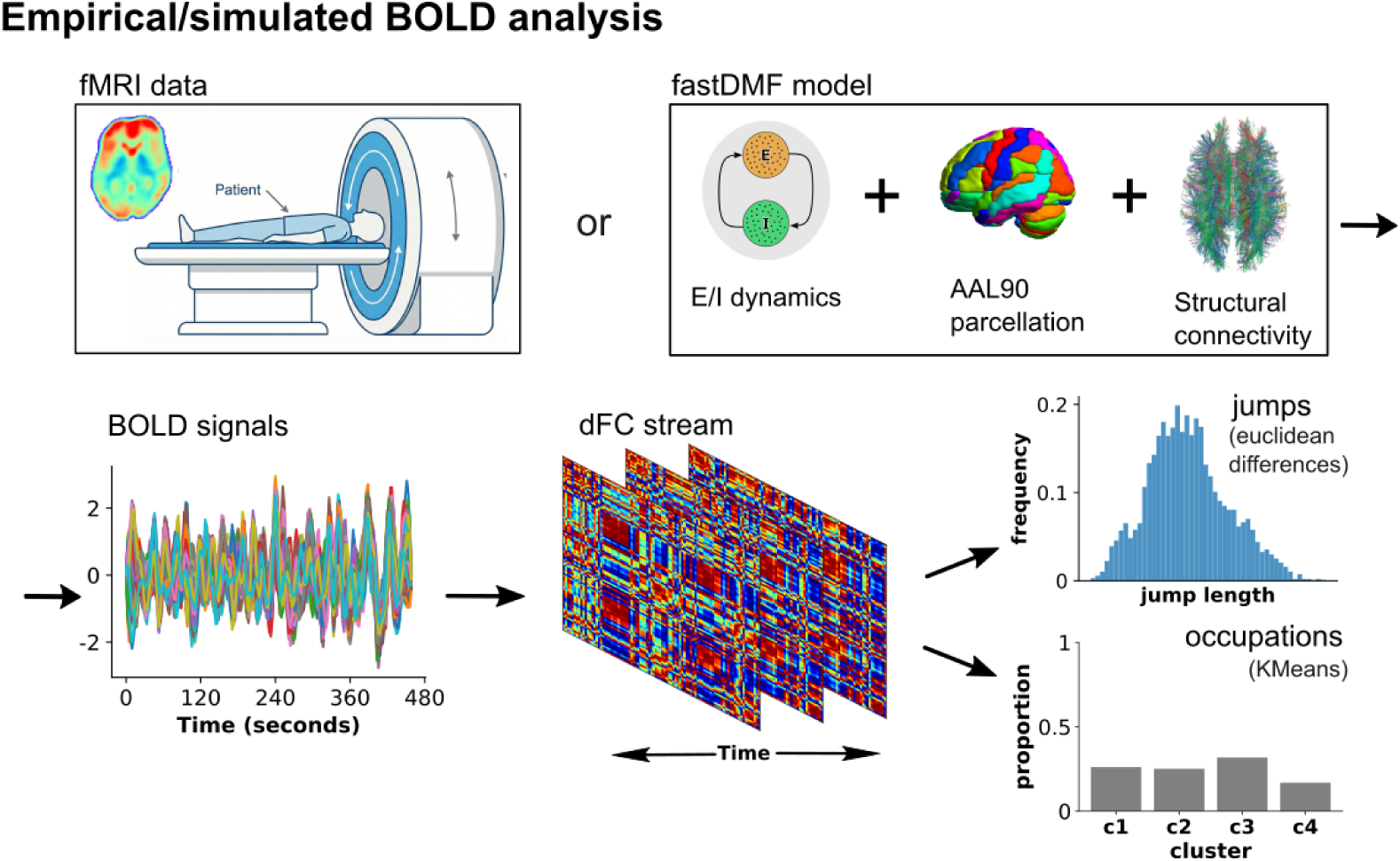
Data analysis pipeline. Empirical fMRI or simulated BOLD (blood-oxygen-level-dependent) signals from the fast dynamic mean-field model (fastDMF), coupled through the structural connectivity (SC) of the 90-region Automated Anatomical Labeling (AAL90) parcellation, were band-pass filtered (0.01-0.1 Hz) and converted into instantaneous phases with the Hilbert transform. At each time point, the phase coherence between every pair of regions gives a dynamic functional connectivity (dFC) matrix, and the sequence of matrices forms the dFC stream. Two observables are computed from it: jump lengths, the Euclidean distance between consecutive dFC matrices, and cluster occupancies, the proportion of frames assigned to each of four recurrent connectivity states by K-means clustering.

## Results

### dFC stream jump length distribution

We first sought to characterize the temporal dynamics of FC changes. For this, we calculated the instantaneous phases of BOLD signals from regions defined by the AAL parcellation [26, 27]. This allowed us to represent each time point with a FC matrix of phase connectivity (see Methods), which we call the dynamic FC stream (dFC stream). We then calculated the euclidean distance between consecutive FC matrices, obtaining a distribution of jump lengths for each recording session. This is what we call “jump lengths” in the dFC stream: larger jump lengths imply bigger changes in the connectivity profile between successive times.

On average, CNT individuals displayed shorter jumps than MCS and UWS (mean jump length 24.6, 26.8 and 27.3), but their jump lengths were more variable (coefficient of variation 0.37, 0.27 and 0.24; Fig 2B). We fitted a Weibull distribution to each individual’s jump lengths (Methods): its scale *λ* sets the characteristic step size and its shape *k* sets the relative spread of step sizes, independently of scale. Fit quality did not differ between groups (Kolmogorov–Smirnov distance, Kruskal–Wallis p = 0.10). CNT jumps had a lower scale (*λ* = 275 vs 29.5 and 29.8; corrected p = 0.042 and 0.036) and a lower shape parameter (*k* = 305 vs 419 and 471; corrected *p* = 0008 and 0003), with no differences between MCS and UWS (Mann–Whitney tests after Kruskal–Wallis, Benjamini–Hochberg corrected). A lower *k* means a broader distribution relative to its typical value, with relatively more very short jumps and relatively larger leaps; each individual’s k predicted their empirical tail ratios (median/5th percentile: *r* = 088; 95th percentile/median: *r* = 097). Jump lengths were also more persistent in time in CNT (lag-1 autocorrelation 0.65 vs 0.53 and 0.55; corrected *p* = 0018 and 0.007), indicating that controls alternate between episodes of slow and fast change in FC.

**Fig 2.**
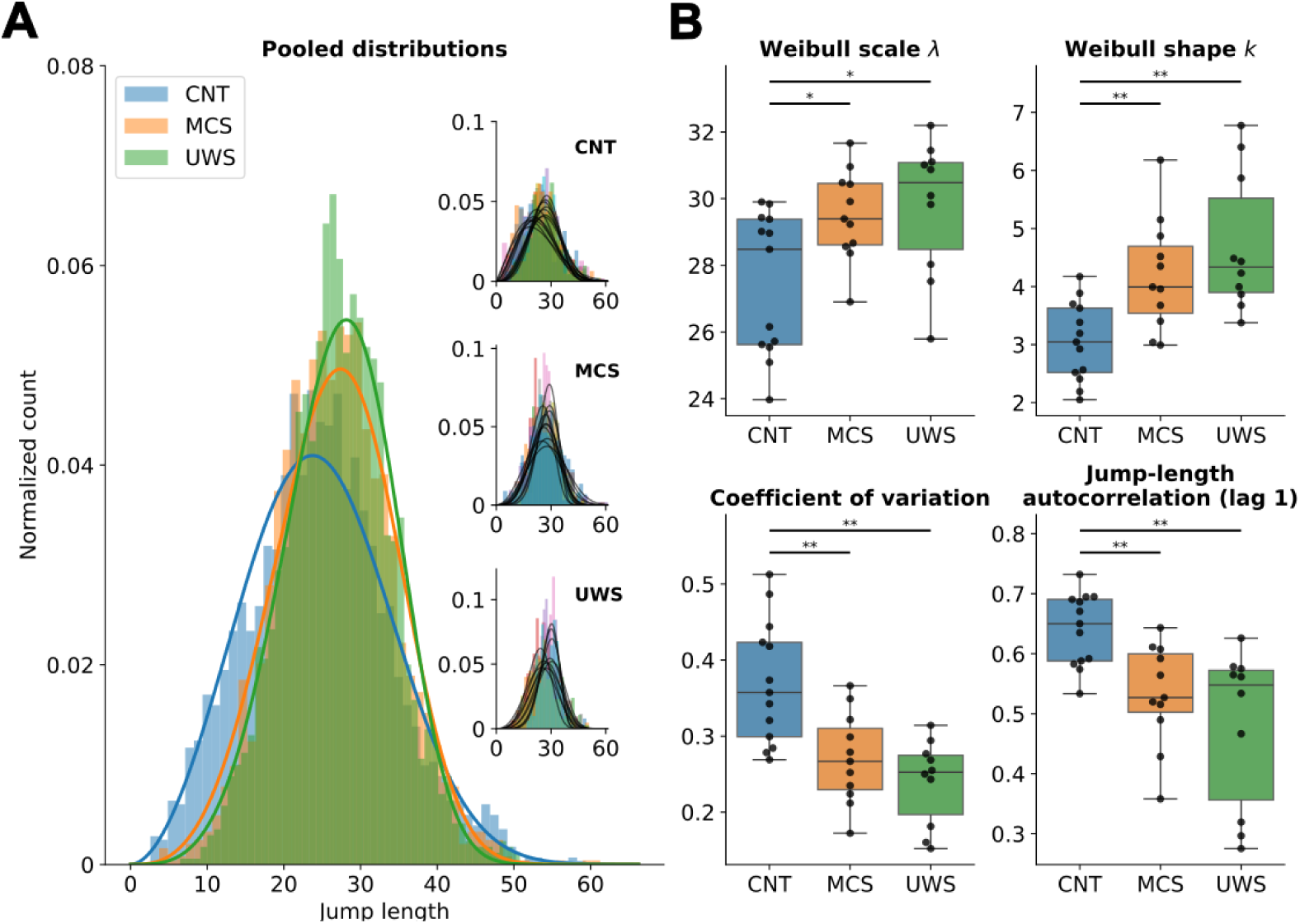
Jump-length distributions of empirical dFC streams. (A) Pooled jump-length distributions of healthy controls (CNT, *n* = 13), minimally conscious state (MCS, *n* = 11) and unresponsive wakefulness syndrome (UWS, *n* = 10) patients, with Weibull fits. Insets show each individual’s distribution and fit (black lines). (B) Per-individual Weibull scale *λ* and shape *k*, coefficient of variation, and lag-1 autocorrelation of jump lengths. The coefficient of variation is computed directly from the jump lengths, without assuming a distribution; under the Weibull model it decreases with *k*, so its agreement with *k* shows that the group differences in relative spread do not depend on the fit. Kruskal-Wallis tests across groups (*λ*, *p* = 0016; *k*, *p <* 0001; coefficient of variation, *p <* 0001; autocorrelation, *p* = 0001) were followed by pairwise Mann-Whitney tests (Benjamini-Hochberg corrected): * *p <* 005, ** *p <* 001, *** *p <* 0001.

### Occupancy of dynamic states of connectivity in CNT and DoC

After characterizing temporal changes in the dFC stream, we studied the main spatio-temporal patterns governing brain activity. For this, we leveraged on a cluster-based analysis developed in [4], and reproduced in several other studies [28] [29] [30]. We used a K-Means clustering algorithm (K=4, see Methods), trained over all concatenated and flattened time-resolved FC matrices of the dFC stream. We sorted centroids according to their resemblance to SC, as measured by the FC-SC correlation coefficient of each centroid (see Fig 3A).

**Fig 3.**
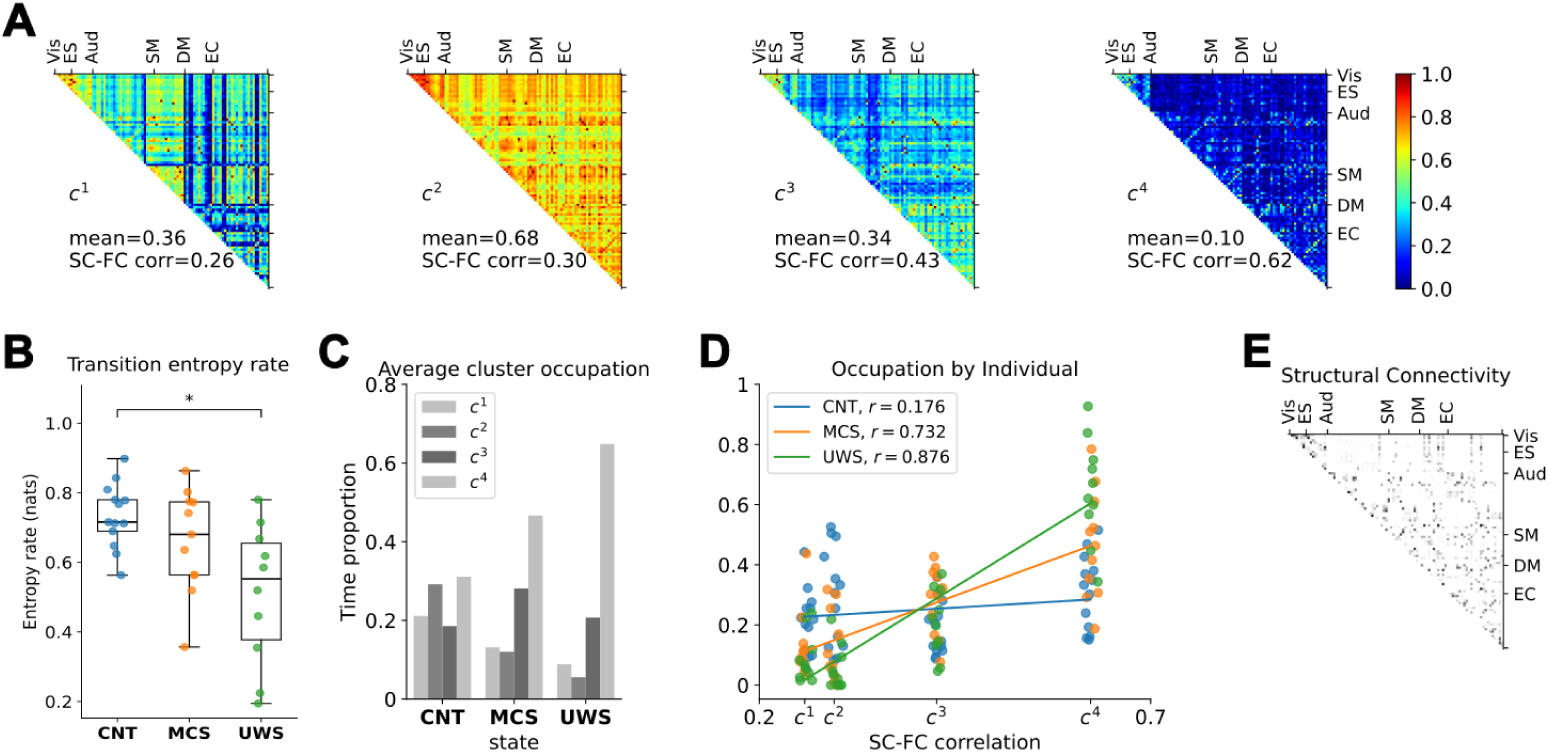
Recurrent dFC states and their occupancy. (A) Centroids of the four K-means clusters (*c*^1^-*c*^4^), ordered by their correlation with the SC matrix (SC-FC correlation). Regions are grouped by resting-state network: visual (Vis), extrastriate (ES), auditory (Aud), sensorimotor (SM), default mode (DM) and executive control (EC). Each panel gives the centroid’s mean coherence and SC-FC correlation. (B) Entropy rate of each individual’s sequence of clusters (first-order Markov chain; Methods). Kruskal–Wallis test followed by pairwise Mann–Whitney tests (Benjamini–Hochberg corrected): * *p <* 005. (C) Mean occupancy of each cluster per group. (D) Occupancy of each cluster in each individual against the cluster’s SC-FC correlation, with a linear fit and Pearson r per group (points jittered horizontally for visibility). (E) SC matrix, ordered as in A.

From the algorithm’s labels (indicating which cluster the dFC belongs to at each time step), we calculated the transition rate between clusters (Fig 3 B) for CNT, MCS and UWS, from which the Markov chain Shannon entropy rate can be estimated. Computed per individual, the entropy rate decreased from CNT to MCS to UWS (0.73, 0.66 and 0.51 nats, respectively; Kruskal–Wallis p = 0.020), with only CNT and UWS differing significantly (corrected p = 0.021).

We define occupancy as the fraction of how many fMRI volumes our K-means algorithm assigned to each cluster, and Fig3 C shows the across-individuals average occupancy of each cluster. We also analyzed occupancies per individual, placing each centroid SC-FC correlation value on an x-axis, and considering the occupancy of each cluster as a function of its FC-SC resemblance. CNT individuals occupied the four clusters at similar rates regardless of their SC–FC similarity (Pearson *r* = 018, *p* = 021), whereas MCS and UWS patients increasingly occupied clusters with higher SC–FC similarity (*r* = 073 and 088, both *p <* 0001; Fig 3D), as previously reported [4]. Fitting one slope per individual confirmed this without pooling points from the same subject: slopes were close to zero in CNT (median 0.13, Wilcoxon *p* = 041) and positive in MCS (1.22, *p* = 0003) and UWS (1.65, *p* = 0002), and differed between all three groups (Kruskal–Wallis *p <* 0001; corrected *p* = 0012, *p <* 0001 and *p* = 0045 for CNT–MCS, CNT–UWS and MCS–UWS).

### Calculating jumps and cluster occupancies using a whole brain model

Next, we assessed whether certain brain regions are more likely to induce changes in the dFC stream. For this, we used a whole-brain model of spontaneous activity fitted to the distribution of jumps and occupancy of connectivity profiles. We generated BOLD-like activity using the fast DMF model [31], over which we calculated the simulated dFC stream using the same pipeline as in empirical data (see Fig 1).

Instead of the widespread strategy of fitting the model to reproduce the FC matrix, we fitted the global coupling parameter *G* aiming to reproduce the empirical jump lengths and occupancies in CNT, MCS and UWS. Jumps were obtained from simulated dFC in the same way as with empirical data. To obtain simulated occupancies, we use the K-Means model that was trained on empirical dFC streams, evaluating its predicted labels on simulated dFC streams. We compared simulated and empirical jump lengths using the Kolmogorov Smirnov distance (KS-distance) on their distributions, and cluster occupancies using the Kullback-Leibler divergence (KL-divergence).

The *G* parameter, that controls the overall strength of interaction between regions, was swept in the *G ∈* [17, 26] range, with a step size of 0.02, and running 20 random seeds for each *G* value. Since KL and KS are dissimilarity measures, the optimal *G* value for each state was chosen as the one that minimizes the distance between empirical and simulated per-state pooled jump lengths (and per-state average occupancies). For jump length distributions and cluster occupancies, the optimal G value decreased monotonically between CNT, MCS, and UWS, suggesting that healthy controls require stronger inter-regional interactions than patients with disorders of consciousness, (jump length optima at *G* = 24, 206, 194 and occupancy optima at *G* = 248, 236, 22 for CNT, MCS and UWS, respectively, see Fig 4A and B).

**Fig 4.**
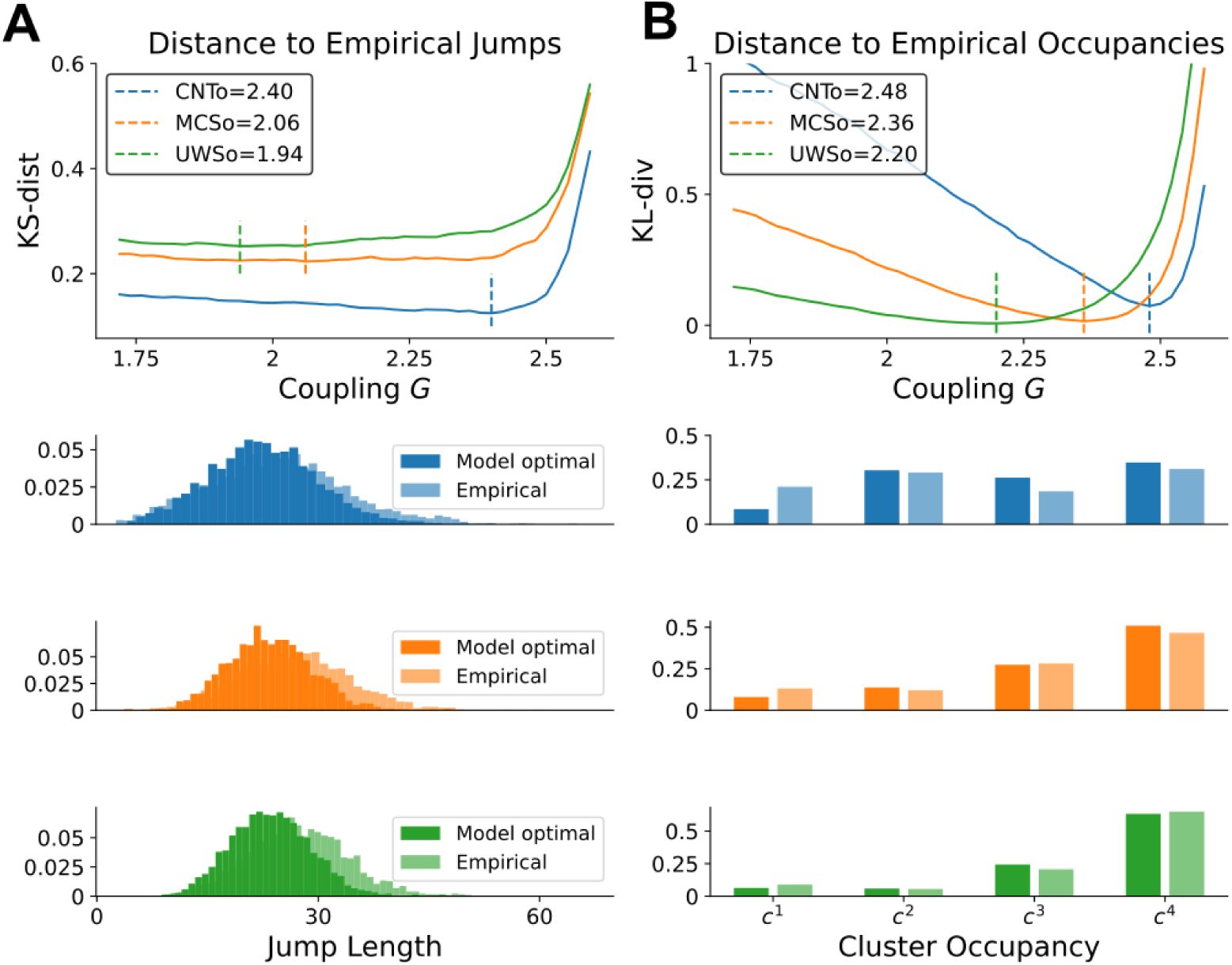
Fitting the whole-brain model to empirical jump lengths and cluster occupancies. (A) Top: Kolmogorov-Smirnov (KS) distance between simulated and empirical jump-length distributions as a function of global coupling *G*, for each group; dashed lines mark the optimal *G* (CNT 2.40, MCS 2.06, UWS 1.94). Bottom: jump-length distributions of the model at each group’s optimal *G* (darker) and empirical distributions (lighter). (B) Same for cluster occupancies, using the symmetrized Kullback-Leibler (KL) divergence (optimal *G*: CNT 2.48, MCS 2.36, UWS 2.20); bottom panels compare simulated (darker) and empirical (lighter) occupancies of *c*^1^-*c*^4^. Each *G* value was simulated with 20 random seeds.

### Nodal integration vs the effect of removing areas from the network

Having obtained the 1) jump length and 2) occupancy optimal *G* values for the CNT condition, we used them as a starting point from which to test the effect of removing each brain area, carrying out the following experiment: Fixing *G* at the CNT optimal, we disconnected each area from the rest of the network (setting the nth row and column of the SC matrix to 0) and re-ran the model, obtaining a “virtual lesion”. We compared its jumps and occupancies to the empirical counterparts of CNT, MCS, and UWS.

We found that for most brain areas, their removal does not significantly affect the ability of model to fit the empirical jump distributions or state occupancy of CNT. However, removing any of ten specific nodes has a much larger impact, making the model to depart from the CNT fit and, importantly, making it closer to the MCS and UWS conditions (y axis in Fig 5 A). Considering that the only heterogeneity between brain areas in our model is their connectivity structure, we sought to look for structural determinants that might explain this fact. We center our analyses on Nodal Integration, that is calculated from a hierarchical SVD decomposition of the SC matrix, represents how globally coupled the system is (see Methods) [25], and is closely related to eigenvector centrality. Calculating this measure for each node sets a clear boundary between low and high integration nodes (that we highlight in Fig 5 B). Removing high integration nodes significantly worsens the fit to CNT jumps when compared against removing low integration ones, and the opposite is true for MCS and UWS (Mann–Whitney, Benjamini–Hochberg corrected *p* = 0035*, d* = 073 for CNT; *p <* 0001 for MCS and UWS, see boxplots in Fig 5 A). Notably, removing the high integration areas can improve the jump length fit to MCS even beyond its own *G*-sweep optima (compare with baseline in Fig 5A). Removing high-integration nodes also made the model’s jumps less variable, raising the Weibull shape parameter from 324 in the intact model to a median of 370 (other nodes: 334; Mann–Whitney *p <* 0001; Fig S1B), toward the values of MCS and UWS patients.

**Fig 5.**
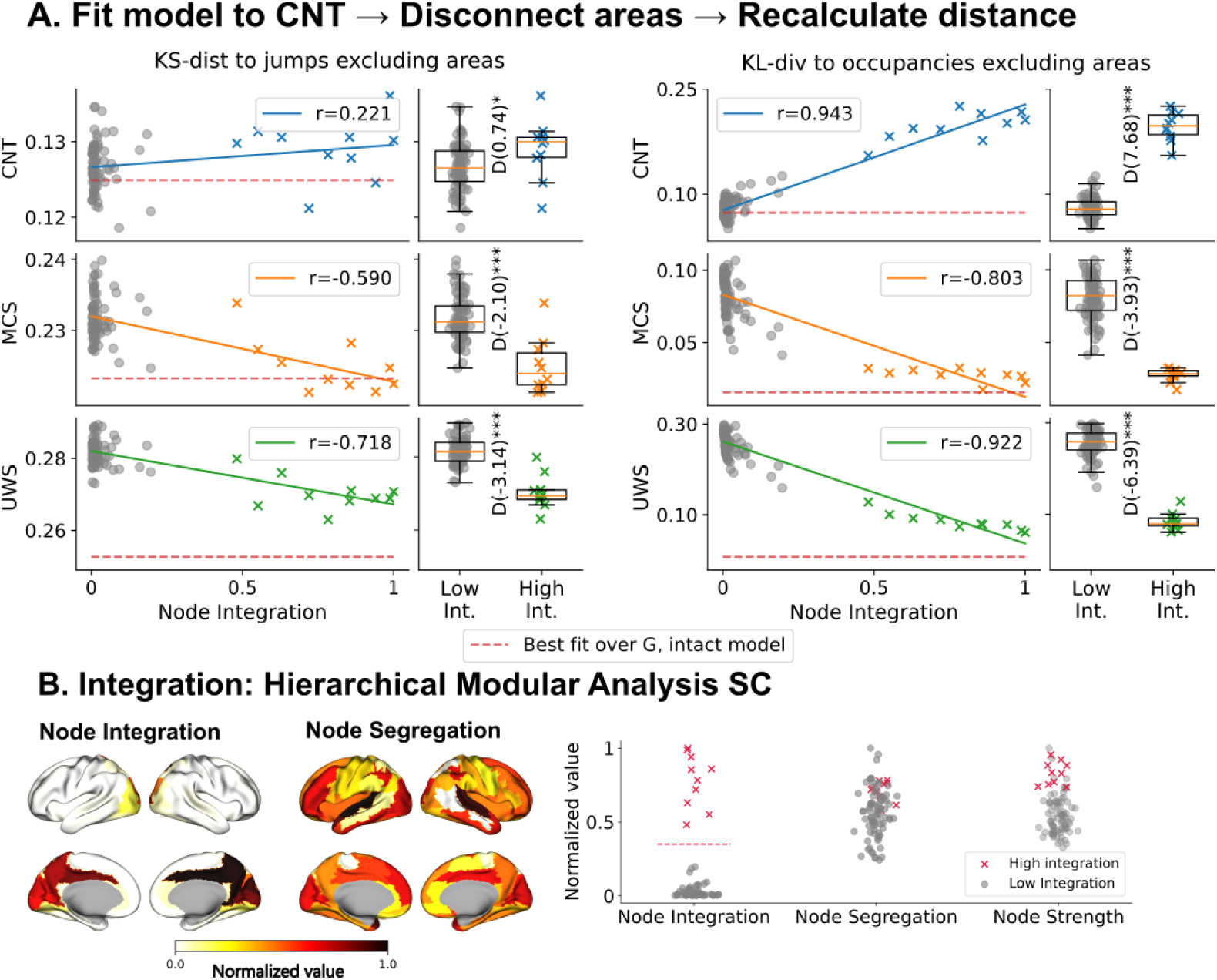
Removing high-integration nodes moves the healthy-fitted model toward patient-like dynamics. Starting from the CNT-optimal coupling (*G* = 240 for jumps, *G* = 248 for occupancies), each node was disconnected by setting its row and column of the SC matrix to zero, and the model was re-simulated with 20 seeds. (A) Left: KS distance between the lesioned model’s jump lengths and the empirical jump lengths of CNT, MCS and UWS (rows), against the normalized integration of the removed node (grey circles: other nodes; colored crosses: the 10 high-integration nodes), with Pearson *r*. Red dashed lines mark each group’s best fit in the *G* sweep (Fig 4). Boxplots compare removing low-versus high-integration nodes (Mann-Whitney tests, Benjamini-Hochberg corrected; D, Cohen’s *d*). Right: same for the KL divergence to each group’s cluster occupancies. (B) Integration and segregation of each node from the hierarchical modular analysis of the SC matrix, shown on the cortical surface, and normalized integration, segregation and strength of all nodes. Integration is bimodal: any threshold between 0.20 and 0.48 selects the same 10 high-integration nodes (posterior cingulate, precuneus, cuneus, superior occipital and calcarine cortex, all bilateral).

For occupancies, removing any of the high-integration areas worsens the fit to CNT and makes the model’s occupancies be more similar to the ones of MCS and UWS, as measured by a decrease in the KL-divergence to each states’ occupancies, however not as close as their own optimal fit from the *G* sweep (see baseline lines in Fig 5A). Removing high integration nodes significantly worsens the fit to CNT compared to low integration nodes, and the opposite is true for MCS and UWS occupancies (*p <* 0001 for CNT, MCS and UWS, see brain areas in Fig 5B). Across nodes, integration correlates more strongly than node strength with the lesion effect on occupancies relative to CNT (Pearson *r* = 094 vs 058) and on jumps relative to MCS and UWS (*r* = *−*059 vs *−*023 and *−*072 vs *−*038; all corrected *p <* 0001), whereas both measures performed similarly for occupancies relative to MCS (*r* = *−*080 vs *−*087; Table S1).

### Stimulating Brain Areas

In the previous section we performed structural “virtual lesions” to a model fitted to CNT dynamics comparing it to DoC observables. Here, we assess which nodes can be functionally modulated to shift a model fitted to DoC back towards healthy dynamics. For this, we perturbed the E/I balance of each node by sweeping the local inhibitory feedback parameter intercept *C* of the fast DMF model, a parameter that increases local inhibition (see Methods). We set our baseline as 1) the best fit to UWS jump lengths (*G* = 194) and 2) the best fit to UWS empirical occupancies (*G* = 22), and compared the perturbed model’s fit to CNT jumps and occupancies. The optimal fits to CNT empirical jumps and occupancies are found with negative *C* values, which are to be interpreted as higher excitability of each node that is required to approximate CNT dynamics.

For each node we took its best fit over the C sweep, restricted to C values for which *J_F IC_* remains non-negative (Methods); without this restriction, the best fits of 28 nodes (jumps) and 58 nodes (occupancies) fell where *J_F IC_ <* 0, none of them high-integration nodes. Stimulating high-integration nodes restored the fit to CNT jumps and occupancies far better than stimulating other nodes (best KS distance, *d* = 277; best KL divergence, *d* = 244; both *p <* 0001, Mann–Whitney; see Fig6). High-integration nodes reached their best fits at *C* between *−*20 and *−*10 for jumps and between *−*08 and *−*06 for occupancies. Stimulating them also made the UWS model’s pace more variable, lowering the Weibull shape parameter from 467 to a median of 410 (other nodes: 436; Fig S1C). The single best jump fit came from stimulating the left middle occipital gyrus, the most integrated node outside the core, at near-complete loss of local feedback inhibition.

### Integration and other centrality measures

Nodal integration ranks nodes in the same order as eigenvector centrality. The separation of ten high-integration nodes reflects the localization of the leading eigenvector of the structural connectome. No other centrality measure predicted the effects of lesions or stimulation consistently better: node strength, and betweenness for CNT jumps, reached similar or slightly higher correlations for three effects (lesion effect on CNT jumps and MCS occupancies, stimulation effect on occupancies), without significant differences (Table S1). For stimulation, the best fit correlated with integration (Pearson *r* = *−*070 for jumps and *−*066 for occupancies) and similarly with node strength (*r* = *−*057 and *−*066; difference not significant), but within the 80 nodes outside the core, integration tracked the effect of stimulation on jumps far better than strength (*r* = *−*061 vs *−*031). The decisive feature is therefore not a general superiority of integration over strength, but the sharp boundary that integration draws: high-integration nodes had larger effects than the remaining nodes in all eight comparisons, six for lesions and two for stimulation, with Cohen’s d of at least 2.1 in seven of them (Figs 5 and 6).

**Fig 6.**
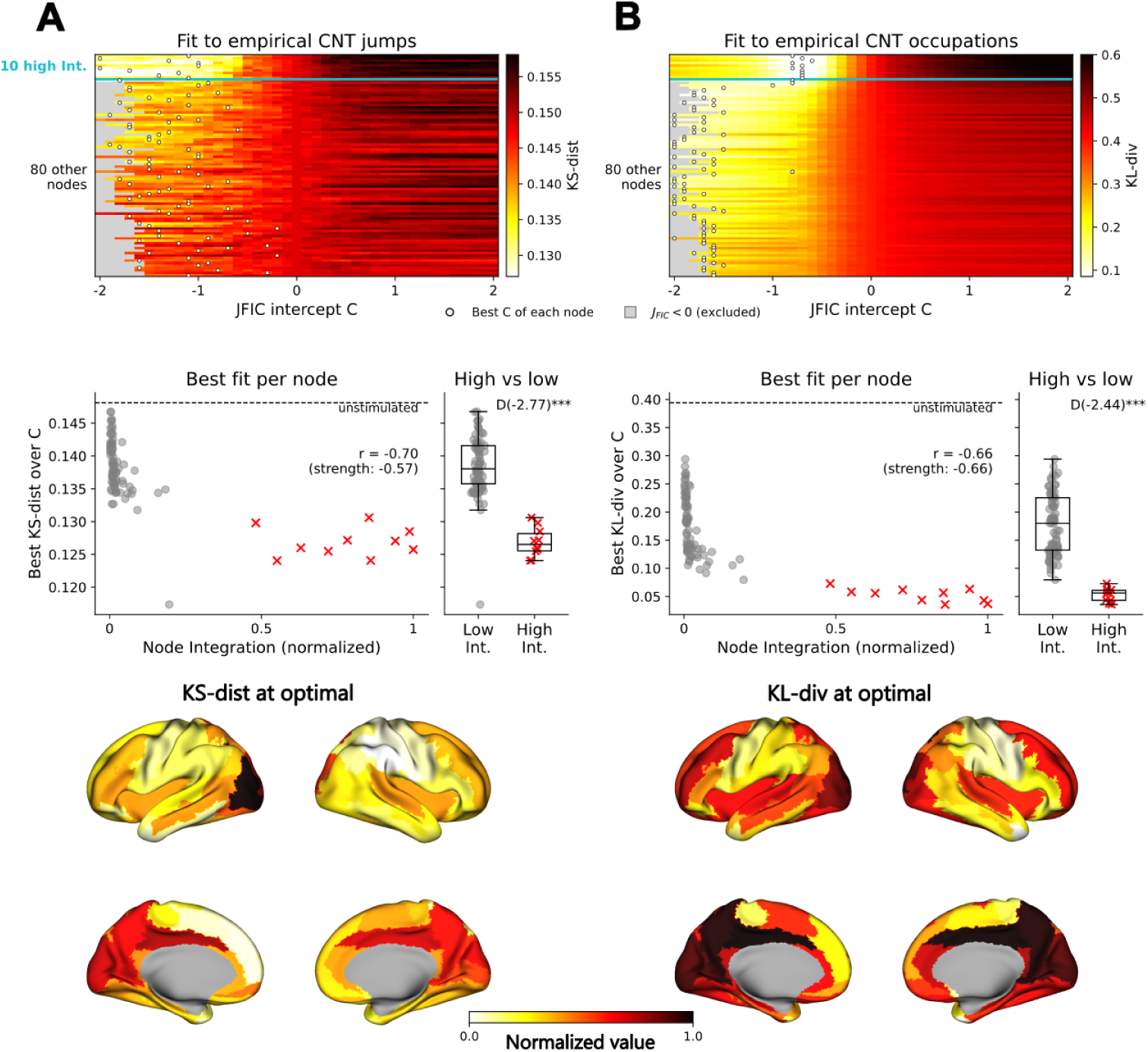
Stimulating high-integration nodes moves the UWS-fitted model toward healthy-like dynamics. Starting from the UWS-optimal coupling, the excitability of each node was changed by shifting the intercept *C* of its feedback inhibition control, *J_F IC_*(lower *C*, higher excitability), and the fit to the empirical CNT data was recomputed. (A) Fit to CNT jump lengths (KS distance; *G* = 194). (B) Fit to CNT cluster occupancies (KL divergence; *G* = 220). Top: fit for each node (rows, sorted by integration; the cyan line separates the 10 high-integration nodes) and each value of *C*. Grey cells mark values of *C* for which *J_F IC_ <* 0, which were excluded; white dots mark each node’s best *C*. Middle: each node’s best fit over the valid range of *C* against its normalized integration (red crosses: high-integration nodes; dashed line: unstimulated model), with Pearson *r* for integration and for node strength; boxplots compare low-versus high-integration nodes (Mann-Whitney test; D, Cohen’s *d*). Bottom: best fit of each region on the cortical surface, normalized between 0 and 1.

## Discussion

In this study, we examined how large-scale structural connectivity constrains the sensitivity of whole-brain dynamics to perturbations, focusing on transitions between dFC regimes characteristic of CNT and DoC conditions. First, we quantified dynamical richness using measures derived from transitions in functional connectivity space. Using perturbations on a whole-brain model, we found that applying the Hierarchical Modular Analysis integration component to the SC matrix reveals nodes that are central to large-scale brain connectivity, as regions with highest integration component best explain transitions from CNT-like to DoC-like dynamics when removed from the network, and transitions from DoC-like to CNT-like dynamics when perturbed toward higher excitability.

In controls, jump lengths were broader relative to their typical size (lower Weibull shape parameter) and more persistent in time (higher lag-1 autocorrelation), indicating that healthy FC alternates between episodes of slow and fast reconfiguration. In patients, jump sizes were more uniform and less correlated in time. This is consistent with the view that conscious brain dynamics are metastable, alternating between transient stability and rapid reconfiguration [32]. In the model, removing high-integration nodes raised the shape parameter toward patient values, and exciting them lowered it toward control values (Fig S1), linking this dynamical signature to the structural core.

Our integration measure ranks nodes exactly as eigenvector centrality, which Váša et al. [23] identified, together with connector hubs, as the best predictor of how a lesion alters synchrony and metastability in a Kuramoto model. Our results are consistent with theirs in a different model class and extend them in two ways: the observables are fitted to empirical dFC in controls and patients, and the same nodes drive transitions in both directions, toward DoC-like dynamics when removed and back toward healthy-like dynamics when excited. The sharp separation of ten high-integration nodes reflects the localization of the leading eigenvector of the structural connectome on the posterior medial structural core described in [16]. On the contrary, nodal segregation did not predict perturbation effects.

In [20], the authors also forced transitions out of DoC-like regimes in silico, stimulating each region of a Hopf model fitted to the state probabilities of an independent cohort. Pushing single regions into oscillation moved both patient models toward controls, with the somatomotor network, thalamus, precuneus and insula as the most sensitive targets, and sensitivity related to how closely each region’s fitted effective connectivity matched healthy structural connectivity. Our results agree on the precuneus and differ in three respects: we changed local excitability through feedback inhibition rather than the oscillatory regime, we perturbed in both directions, and we kept the normative connectome fixed rather than fitting effective connectivity per group. Keeping the connectome fixed isolates the contribution of each node’s position in the structural network, and under these conditions a node’s effect, whether removed or excited, followed its integration component.

Highly integrated nodes coincide with the posterior medial structural core described by [16], and only partly overlap with the rich club [33]: rich-club regions such as the superior frontal gyrus and putamen have high strength but low integration in our connectome, so integration reflects membership in the dominant structural mode rather than degree alone. While the precise anatomical identity of such hubs can depend on methodological choices such as parcellation and binarization thresholds [33], our results consistently identify the posterior medial core, led by the posterior cingulate and precuneus in occupancy-based measures. The relevance of the Precuneus for consciousness-related processes is well established [34], and its activity is consistently reduced in DoC patients [35], while its connectivity profile has been shown to predict clinical outcomes following coma [13]. A methodological advantage of the integration measure used here is that it does not depend on arbitrary thresholding of the SC matrix. The Hierarchical Modular Analysis framework was introduced on the structural connectome [24] and later applied to functional connectivity [25]; being defined over a spectral decomposition, it applies to any symmetric connectivity matrix.

SC sets the conditions under which FC emerges [36]. From a dynamical systems perspective, highly integrative hubs occupy bottleneck positions in state space, such that local perturbations disproportionately affect global trajectories. Identifying which structural characteristics best predict the controllability of a network toward a desired working point is a question of broad interest in mathematics and biology. In neuroscience, this perspective is particularly relevant to understand how targeted interventions can modulate pathological brain dynamics. Non-invasive stimulation techniques such as transcranial magnetic stimulation (TMS) have shown measurable improvements in clinical scales on DoC patients when applied to regions such as the dorsolateral prefrontal cortex [37] or posterior parietal areas [38] [39]. Our results provide a structural rationale for why perturbing integrative hubs may be particularly effective in reshaping global brain dynamics.

Consistent with this interpretation, shifting the local excitation–inhibition balance toward excitation in high-integration regions in our model is sufficient to restore healthy-like dynamics from a DoC regime. This finding aligns with the view that DoC corresponds to a network operating in a sub-excitable state [40] [35], in which activity fails to propagate and integrate across cortical hubs [41].

The thalamus did not emerge as a highly influential node in our simulations, despite prior work identifying it as crucial for sustaining arousal, using a similar pipeline for estimating dFC [42], and despite in silico stimulation identifying thalamic segments among the effective targets for DoC [20]. On the contrary, a node’s influence in our framework follows its weight in the leading eigenvector of the SC matrix, which is concentrated on the posterior medial cortex. Two methodological differences may explain this discrepancy: in [20], the authors work using a Hopf model, in which the parcellation used divides the thalamus into segments with distinct cortical projections, whereas the AAL atlas represents each thalamic hemisphere as a single region that averages nuclei with different targets, and they fitted effective connectivity to each group’s FC, which can strengthen thalamocortical couplings beyond those estimated by tractography. More generally, neither model captures the cytoarchitecture of thalamic nuclei or the full complexity of corticothalamic loops [43].

An interesting observation is that DoC patients tend to occupy connectivity clusters with higher SC-FC similarity, yet lower global coupling values are required to fit DoC dynamics than CNT dynamics. Similar findings have been reported when fitting different fMRI-derived observables [18] [44] [17]. We interpret this as reflecting the capacity of the healthy human connectome to support emergent dynamics that are constrained by structure [19], yet dynamically richer. Further work using more detailed models will be necessary to investigate this phenomenon in greater depth.

Several limitations should be acknowledged. Choosing a different parcellation may partially alter the anatomical identity of which nodes are highly integrated (preserving the approximate position on the cortex), although preserving the underlying spectral and topological features that give rise to global integration. We expect the observed relationship between nodal integration and dynamical impact to be robust to changes in parcellation, even though the specific nodes identified as hubs may vary. In addition, the whole-brain model relies on simplifying assumptions, including predominantly excitatory interareal coupling and a fixed hemodynamic response model, which are expected to affect quantitative aspects of the dynamics rather than the qualitative relationships reported here. Stimulation was implemented as a simplified modulation of local E/I balance, providing a controlled and interpretable perturbation but not capturing the full biophysical complexity of neuromodulatory interventions. A further limitation is that we relied on a normative structural connectome, which may not accurately reflect individual or pathology-specific connectivity in DoC patients. Accordingly, our results are not intended as patient-specific predictions, but rather as a characterization of general relationships between structural integration and dynamical sensitivity that can inform future studies using individualized connectomes. Jump lengths are frame-to-frame differences, so head motion and lower signal quality in patients could contribute to their narrower and less persistent distributions. The sample was small (13, 11 and 10 individuals) and models were fitted at the group level. Finally, the best jump fits required strong reductions of local feedback inhibition, and all nodes were stimulated over the same range of *C* despite different baseline feedback inhibition.

Taken together, our results show that generic features of large-scale structural connectivity, as measured by diffusion MRI, constrain the sensitivity to perturbation of whole-brain dynamical states and their transitions between brain-wide regimes, providing a principled framework for understanding how structural integration shapes sensitivity to perturbation in health and disorders of consciousness.

## Materials and methods

### Participants

We analysed resting-state fMRI data from 11 patients in a minimally conscious state (MCS; 5 women, age 473 *±* 208 years), 10 patients with unresponsive wakefulness syndrome (UWS; 4 women, 393 *±* 163 years) and 13 healthy controls (CNT; 7 women, 425 *±* 136 years), a dataset previously described in [18]. Trained clinicians assessed each patient’s level of consciousness with the Coma Recovery Scale-Revised (CRS-R) [45]. Patients showing behaviours that may indicate awareness, such as visual tracking, localization of noxious stimuli or reproducible responses to commands, were diagnosed as MCS; patients with preserved arousal (eye opening) but no sign of awareness were diagnosed as UWS. The study was approved by the French ethics committee Comité de Protection des Personnes Île de France 1 (Paris, France) under the designation ‘Recherche en soins courants’ (NEURODOC protocol, no. 2013-A01385-40). Informed consent was obtained from the patients’ relatives and from the healthy controls in accordance with the Declaration of Helsinki and French regulations.

### BOLD time series extraction

All participants were scanned with the same protocol on a 3T General Electric Signa system. Resting-state images were acquired with a T2*-weighted gradient-echo echo-planar imaging (EPI) sequence in axial orientation (200 volumes, 48 slices, slice thickness 3 mm, repetition time/echo time (TR/TE) 2400/30 ms, voxel size 34375 *×* 34375 *×* 34375 mm, flip angle 90*^◦^*, field of view 220 mm), giving 8 minutes of data per participant. A T1-weighted magnetization-prepared rapid gradient-echo (MPRAGE) anatomical image was acquired in the same session (154 slices, slice thickness 1.2 mm, TR/TE 7.112/3.084 ms, voxel size 1 *×* 1 *×* 1 mm, flip angle 15*^◦^*).

Preprocessing followed [17] [18] and used the FMRIB Software Library (FSL). The first five volumes were discarded. Images were corrected for head motion (MCFLIRT), skull-stripped (Brain Extraction Tool), spatially smoothed with a 5 mm full-width-at-half-maximum Gaussian kernel, high-pass filtered at 0.01 Hz and registered to Montreal Neurological Institute (MNI) space with a rigid-body transformation. Noise was removed with single-subject independent component analysis (MELODIC) followed by FMRIB’s ICA-based Xnoiseifier (FIX); in patients, components related to lesion artefacts were also regressed out. Regional BOLD time series were then extracted for the 90 cortical and subcortical regions of the Automated Anatomical Labeling (AAL) atlas [26]. Empirical and simulated time series were band-pass filtered between 0.01 and 0.1 Hz with a second-order Bessel filter (SciPy) before further analysis.

### Structural connectivity

The model was coupled through a normative structural connectivity matrix from 16 healthy right-handed adults (5 women, age 248 *±* 25 years), described in [46] and also used in [18]. Diffusion-weighted images were acquired on a 3T Siemens Skyra scanner at Aarhus University, Denmark, with 62 gradient directions at *b* = 1500 s/mm^2^, one *b* = 0 volume every 10 diffusion-weighted volumes, and two acquisitions with opposite phase-encoding directions. After preprocessing with the FMRIB Software Library (FSL) diffusion toolbox, probabilistic tractography (FSL probtrackx) was run between the 90 regions of the AAL atlas, sampling 5000 streamlines from each voxel of a region. The connection strength from region *i* to region *j* was the fraction of streamlines seeded in *i* that reached *j*, normalized by the number of voxels in *i*; connections supported by fewer than five streamlines were set to zero. Because tractography is seed-dependent, the *i → j* and *j → i* values were averaged to obtain a symmetric matrix.

### Dynamic FC calculation

We applied phase-based estimation of time-resolved Functional Connectivity. For each timeseries, we use the Hilbert Transform to get the analytical expression of our signals in the complex plane, whose angle we extract to obtain the instantaneous phase of all areas’ BOLD signals *{φ_i_*(*t*) *∈* [*−π, π*]*, i* = 1, 2*,…,* 90*}*, and measure the phase coherence matrix at time t between areas i and j as

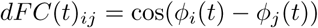

which is equal to 1 when the two areas are in the same phase, zero when they are orthogonal (difference of *π/*2), and-1 if they are in antiphase (difference of *π*). Given that cos is an even function, these matrices are symmetric, so we take the lower triangular entries (4005 in total) to perform our analyses.

### Clustering algorithm on dFC matrices

A K-means algorithm was applied on the concatenated lower triangulars of all individuals, using the scikit-learn native implementation of the algorithm. The number of clusters (K=4) was chosen such that the variability between the inter-cluster correlations is maximized, subject to that the occupancy of each cluster was greater than zero by all three groups.

### Jump length calculation and Weibull fit

Given the flattened lower triangular part of the dFC stream matrices, we calculated the jump length at time t as the euclidean distance between matrices corresponding to consecutive time points *JL*(*t_i_*) = euc(*dFC*(*t_i_*) *− dFC*(*t_i−_*_1_)).

We modelled each individual’s jump lengths with a two-parameter Weibull distribution, *F* (*x*) = 1 *−* exp[*−*(*x/λ*)*^k^*], fitted by maximum likelihood with the location fixed at zero (scipy.stats.weibull_min). Jump lengths are positive, unimodal and right-tailed, and the Weibull family describes them with two interpretable parameters. The scale *λ* sets the characteristic jump length (the 63rd percentile). The shape *k* sets the spread of jump lengths relative to their typical size, independently of scale: the coefficient of variation and every quantile ratio depend on *k* alone, *q_p_/q_r_* = [ln(1 *− p*)*/* ln(1 *− r*)]^1^*^/k^*. A lower *k* therefore means relatively more very short jumps (for small *x*, *P* (*X < x*) *≈* (*x/λ*)*^k^*) and relatively larger ones. Because *k* is estimated from all jumps of an individual, it is a more stable summary than tail quantiles, which rest on few observations. We assessed fit quality with the Kolmogorov-Smirnov distance between each individual’s jump lengths and their fitted distribution.

### Temporal structure of jump lengths and cluster sequences

For each individual, we quantified the variability of jump lengths with their coefficient of variation (standard deviation divided by mean), and their persistence in time with the lag-1 autocorrelation, *ρ*_1_ = corr (*JL*(*t_i_*)*, JL*(*t_i_*_+1_)), the Pearson correlation between consecutive jump lengths. A value of *ρ*_1_ close to zero indicates that each jump is independent of the previous one, whereas higher values indicate that large and small jumps cluster into episodes of fast and slow reconfiguration of FC. Because consecutive phase-coherence matrices share signal through filtering and the Hilbert transform, *ρ*_1_ is positive in all individuals, and we interpret differences between groups rather than its absolute value.

To quantify how predictable transitions between clusters were, we labelled each volume with its cluster and counted transitions *n_ij_* from cluster *i* to cluster *j* between consecutive volumes of the same individual, excluding transitions between the end of one recording and the start of the next. With transition probabilities 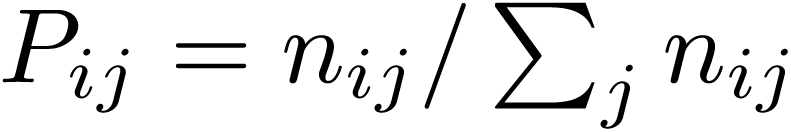 and weights 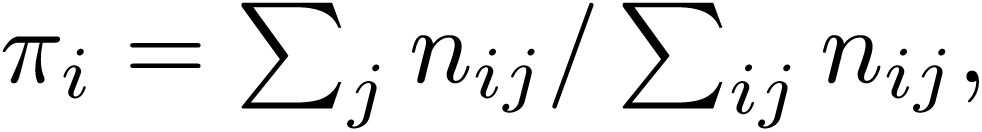 the entropy rate of the corresponding first-order Markov chain is

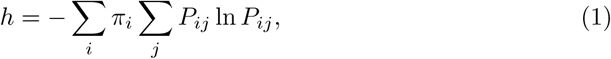

in nats, ranging from 0 (fully predictable transitions) to ln 4 *≈* 1.39 (all clusters equally likely at every step).

Finally, we measured the similarity of each cluster to the structural connectome as the Pearson correlation between the lower-triangular entries of its centroid and those of the SC matrix. To relate cluster occupancies (see *Goodness of fit*) to this similarity without pooling the four non-independent values of each individual, we fitted one least-squares line per individual (four occupancies against the four SC-FC similarities) and kept its slope. Slopes were tested against zero within each group with Wilcoxon signed-rank tests and compared between groups as described in *Effect size estimation*.

### Dynamic Mean Field model (fastDMF implementation)

The Dynamic Mean Field model (DMF) of brain activity [46] is a group of stochastic differential equations that model the dynamics of the interactions between different brain areas. We use the FastDMF implementation in C with a python API, with default parameters [31]. The equations that define the excitatory (*E*) and inhibitory (*I*) dynamics of each node *n ∈ {*1, 2*,…,* 90*}* are:

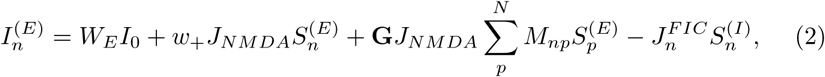

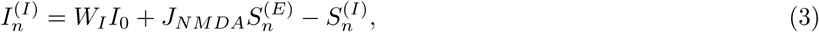

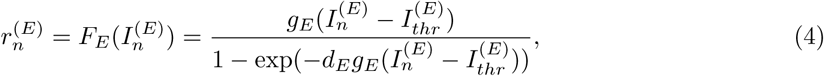

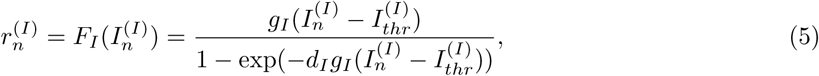

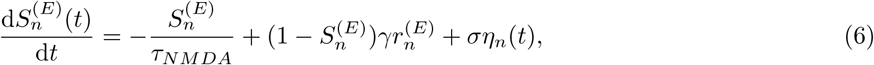

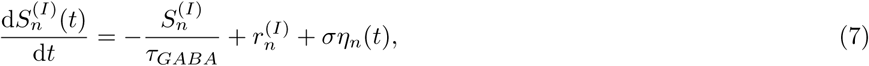

where *I_n_*^(E),(I)^ is total input current, *r_n_*^(E),(I)^ is firing rate, and *S_n_*^(E),(I)^ is the synaptic gating variable of the (*E*) and (*I*) populations. *F_E_*(*·*), *F_I_* (*·*) are transfer functions that define the relationship between input current and output firing rate of the (*E*) and (*I*) populations. As is custom in brain modeling, brain areas are connected through their excitatory activities, according to a weighted undirected SC matrix *M*, whose influence is proportional to the coupling parameter **G**.

*J_n_^FIC^* is the local Feedback Inhibition Control (FIC) parameter, that was shown to optimally follow a linear dependency with node strength [31]:

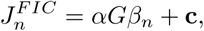

where *β_n_* = ∑*_p_ M_np_* is the node strength of the *n−*th area. In our work, the intercept is subdivided in a baseline and a change parameter **c** = *c* + *C*, with *c* = 1 and a *C* parameter that we change to modulate the local E/I balance. All other parameters are set as in [31].

### Balloon-Windkessel Hemodynamic model of BOLD response

BOLD signals are obtained with an online-calculation (on the run) of the Balloon-Windkessel model [47] [48]. In this model, an increase in excitatory activity induces an exponentially decaying vasodilatory response *s_n_*, which in turn triggers and is affected by blood inflow *f_n_* and changes in blood volume *v_n_* and deoxyhemoglobin content *q_n_*, following:

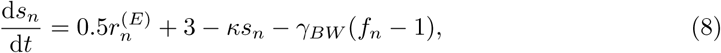

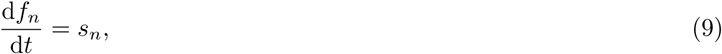

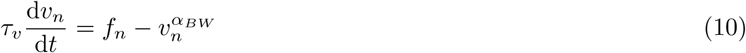

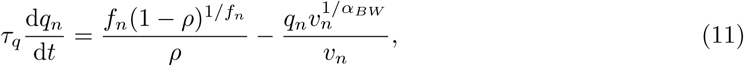

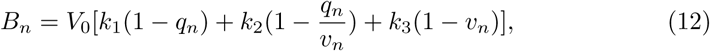

where *B_n_*is the simulated BOLD signal, *ρ* is the resting oxygen extraction ratio, *α_BW_*accounts for mechanical resistance of veins, *τ_q_* is a time constant, and *k*_1_*, k*_2_ and *k*_3_ are coefficients originally estimated from data. All biophysical parameters are the same as in [48].

The model sampling rate was set as *TR* = 24, and simulations had a duration of 8 minutes, from which a transient of 1 minute was discarded from the beginning, in order to work with steady-state dynamics. Filtering was the same as in empirical data.

### Goodness of fit

For comparing empirical and simulated jump distributions, we used the Kolmogorov-Smirnov distance, that we calculate in python using scipy.stats.kstest. This is equal to the greatest absolute gap between two random vectors’ CDFs. Occupancies can be seen as discrete probability distributions, while jump distributions are continuous, so we use different measures for each of them.

To assess the similarity between empirical and simulated occupancies of clusters, we apply the Kullback-Leibler (KL) divergence (equivalent to relative entropy, and calculated in python using scipy.special.rel_entr). Occupancies are defined as the count of fMRI volumes in each cluster (4 values) divided by the total number of volumes, so that they sum up to 1. For obtaining a symmetrical measure, we took the final similarity between probability vectors *{P*_1_*, P*_2_*}* as:

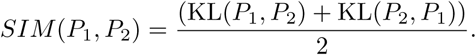

### Effect size estimation

To quantify the magnitude of difference between low-and high-integration node groups (e.g., Fig 5, Fig 6), we used Cohen’s d, computed as:

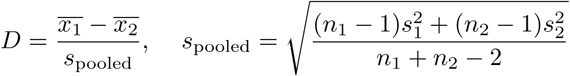

where x̄_1_,x̄_2_ and *s*_1_*, s*_2_ are the means and standard deviations of the two groups to compare, and *n*_1_*, n*_2_ their respective sample sizes. Significance of the difference between groups was assessed using Kruskal Wallis non parametric tests, and then pairwise differences using Mann whitney U-test, with multiple-comparison correction applied via the Benjamini-Hochberg procedure where relevant.

### Integration and Segregation using Hierarchical Modular Analysis

To identify functional modules in the SC matrix, and quantify their integration and segregation components, we used the Hierarchical Modular Analysis (HMA) method [24] [25]. The method applies an SVD decomposition of the SC matrix to find its eigenvectors and eigenvalues (sorted by eigenvalue magnitude, from largest to smallest). The regions whose corresponding entries in a given eigenvector have the same signs are assumed to have joint activity (cooperation) and put in the same module, one for positive and one for negative values. Given that all entries of the SC matrix are positive, the first functional level (largest eigenvalue) consists of a sole one module that encompasses all brain areas, the second level can divide the brain in two modules according to the signs of the entries of the second eigenvector, the third can divide each of these in two more modules, and so on. During this partition process, we save the number of modules in each level *M_i_*, and the size of each module *m_j_*.

The single large module on the first level with the largest eigenvalue Λ_1_ corresponds to global network integration. The second level defines two modules that support local integration within each one and segregation between them, which are weighted by a lower eigenvalue. In subsequent levels, more modules reflect deeper levels of segregation, accompanied by smaller eigenvalues Λ. The weighted module number in each level can be defined to reflect the hierarchical segregated and integrated interactions:

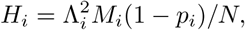

where 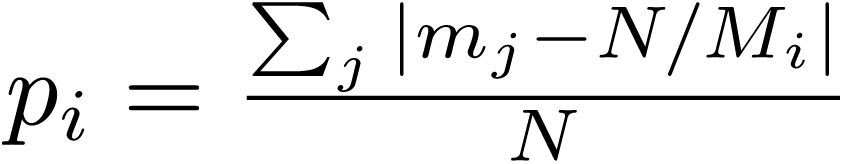 is a correction factor that takes into account heterogeneous modular sizes.

After this, the global integration component is taken from the first level *H*_in_ = *H*_1_*/N*, and the segregation component is obtained as a sum from the 2nd to the *N* th level, as 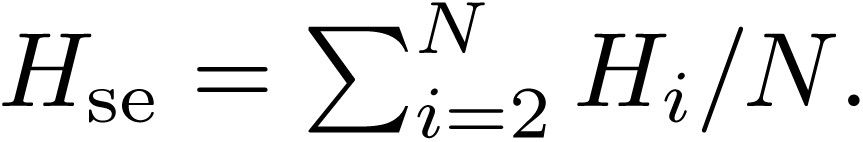 Finally, the local component of segregation *H_se_^j^* and integration *H_in_^j^* for each brain area *j* can be obtained by weighting the integration components for the corresponding entries of the eigenvectors, as:

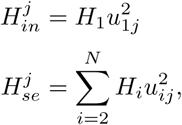

where *u_ij_* is the *j−*th eigenvector entry at the *i−*th level. More details and previous applications of these measures can be found in [24] [25].

Since *H*_1_ is common to all regions and all entries of the leading eigenvector of the SC matrix share the same sign, *H_in_^j^* ranks regions exactly as eigenvector centrality does [49]. Nodal integration therefore measures each region’s contribution to the global integrated mode of the connectome.

## Data Availability

The fMRI data from patients with disorders of consciousness and healthy controls come from a clinical cohort and cannot be made publicly available under the approved ethics protocol; they can be shared by the authors upon reasonable request. The structural connectivity matrix was obtained in previously published work [46]. Simulated data is available at https://zenodo.org/uploads/20842677, and codes needed to reproduce the figures is available at https://github.com/flehue/integration_DoC. Simulations used the FastDMF implementation of the DMF model [31].

## Acknowledgements and Funding

We express gratitude to all patients who participated in the studies, and to their families. F.L. was supported by Beca de Doctorado Nacional N° 21220708 (Agencia Nacional de Investigación y Desarrollo ANID, Chile). I.M. was supported by FLAG-ERA research funding organisation (project ModelDXConsciousness). C. C.-O. is supported by FONDECYT Postdoctoral Fellowship 2026, Agencia Nacional de Investigación y Desarrollo (ANID), Chile, grant 3260021, and by the U-Inicia VID 2026, Universidad de Chile, grant UI-001/26. E. Z. is supported by PICT2019–02294 (I+D+i Agency, Argentina), ANID/FONDECYT Regular 1220995 (Chile), FONDECYT Exploración 13240170 (Chile), and Alzheimer’s Association grant AARG-25-1490306. P.O. is supported by FONDECYT Regular 1241469, Fondecyt Exploración 13240170, the Advanced Center for Electrical and Electronic Engineering (AFB240002 ANID, Chile), and the Instituto Milenio Centro Interdisciplinario de Neurociencia de Valparaíso (ICN2025_026, ANID, Chile). The authors declare no competing interests, and the funding institutions had no role in study design, data collection and analysis, decision to publish, or preparation of the manuscript.

## Notes

https://zenodo.org/records/20842677

https://github.com/flehue/integration_DoC

### Competing Interest Statement

The authors have declared no competing interest.

### Summary of Updates

This version revises the manuscript submitted to Network Neuroscience. We changed the characterization of jump lengths in dFC space, now including a Weibull fit, coefficient of variation and lag-1 autocorrelation, all computed per individual. Statistics now Kruskal-Wallis and Mann-Whitney tests non parametric tests. Stimulation is restricted to parameter values that keep feedback inhibition non-negative. We mention the relation between nodal integration and eigenvector centrality, and include comparisons against other centrality measures. We added supplementary figures and tables and rewrote the Introduction and Discussion in light of prior lesion and stimulation modeling studies. The title was updated.

