## Supplementary figures and tables for "Network integration shapes the loss and recovery of dynamical repertoires in disorders of consciousness"

### Supporting information

Structural determinants of dynamical state transitions in disorders of consciousness: a whole-brain modeling approach

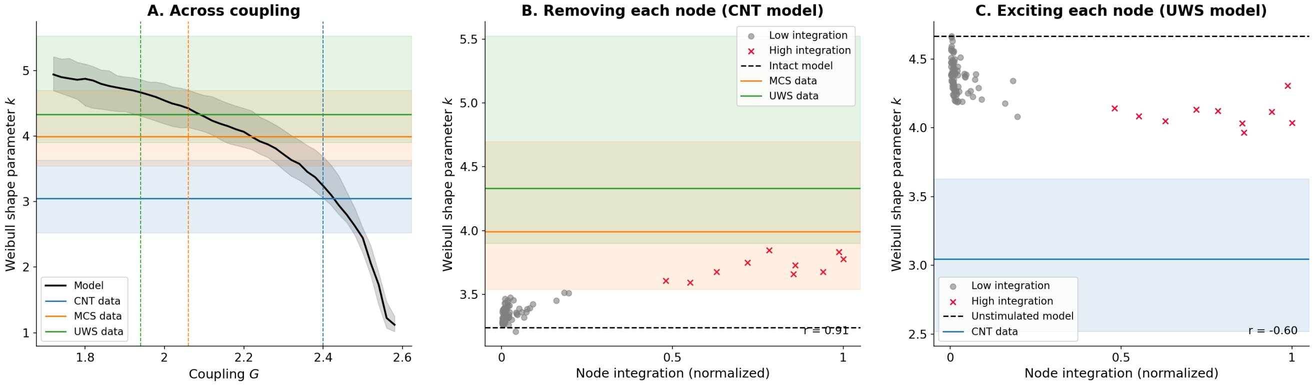


**Fig S1. Weibull shape parameter $k$ of the model's jump lengths. …**

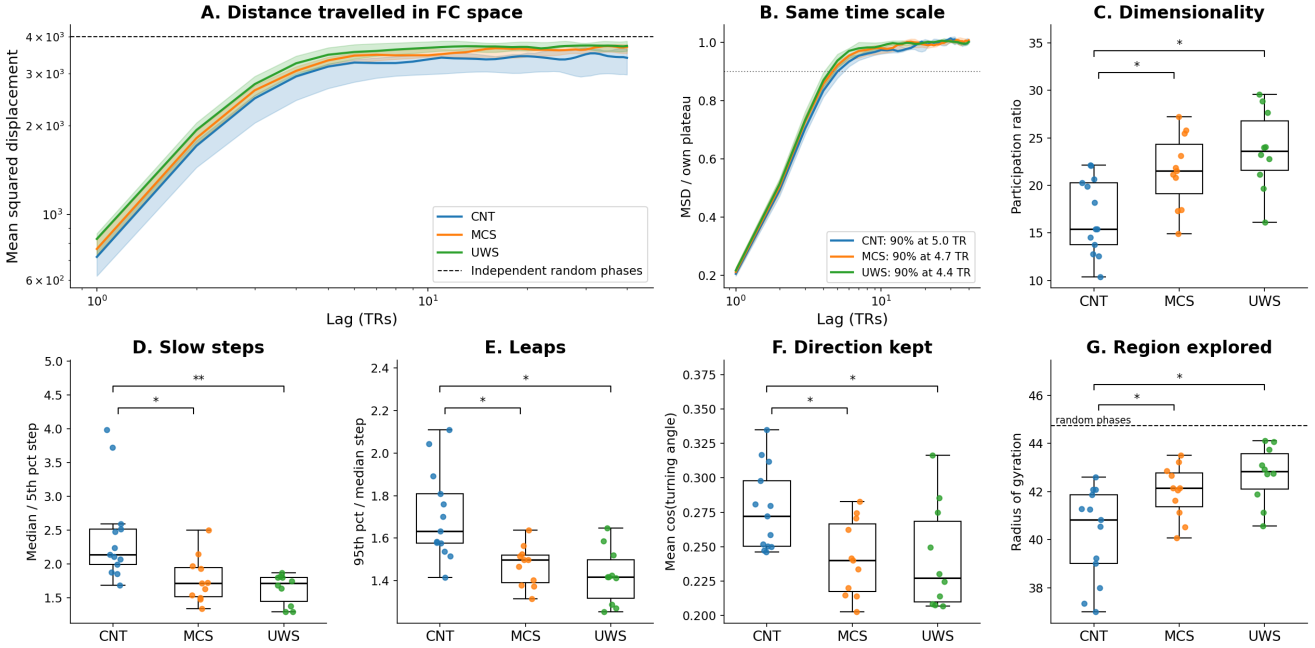


**Fig S2. Exploration of FC space by the empirical dFC streams. …**

**Table S1. Node measures and the effect of perturbing each node.** Correlation across the 90 nodes between each node measure and the effect of removing that node from the CNT-fitted model (lesion: distance to each group's empirical jump lengths, Kolmogorov–Smirnov (KS), and cluster occupancies, Kullback–Leibler (KL)) or of exciting it in the UWS-fitted model (stimulation: best fit to CNT over the values of $C$ with $J_{FIC} \geq 0$). A larger effect means a larger distance to CNT after a lesion and a smaller distance in every other column, so central nodes with large effects give positive correlations in the first column and negative ones elsewhere. Integration ranks nodes exactly as eigenvector centrality, which is not listed separately. Stars mark Benjamini–Hochberg corrected $p$ within each block: $^{*}p < 0.05$, $^{**}p < 0.01$, $^{***}p < 0.001$. The rows marked "outside the core" use only the 80 low-integration nodes. The last Pearson row gives the corrected $p$ of the Meng–Rosenthal–Rubin test comparing the correlations of integration and strength with each effect.

| | Lesion (CNT-fitted model) | | | | | | Stimulation (UWS-fitted model) | |
|---|---|---|---|---|---|---|---|---|
| | Jumps (KS) | | | Occupancies (KL) | | | Jumps (KS) | Occupancies (KL) |
| Node measure | CNT | MCS | UWS | CNT | MCS | UWS | CNT | CNT |
| *Pearson r* | | | | | | | | |
| Integration | 0.22 | $-0.59^{***}$ | $-0.72^{***}$ | $0.94^{***}$ | $-0.80^{***}$ | $-0.92^{***}$ | $-0.70^{***}$ | $-0.66^{***}$ |
| Segregation | $-0.07$ | $-0.16$ | $-0.21$ | $0.29^{**}$ | $-0.41^{***}$ | $-0.38^{***}$ | $-0.19$ | $-0.22$ |
| Strength | $0.28^{*}$ | $-0.23$ | $-0.38^{***}$ | $0.58^{***}$ | $-0.87^{***}$ | $-0.81^{***}$ | $-0.57^{***}$ | $-0.66^{***}$ |
| Betweenness | $0.26^{*}$ | $-0.09$ | $-0.21$ | $0.38^{***}$ | $-0.60^{***}$ | $-0.57^{***}$ | $-0.39^{***}$ | $-0.55^{***}$ |
| Closeness | 0.20 | $-0.07$ | $-0.20$ | $0.39^{***}$ | $-0.68^{***}$ | $-0.63^{***}$ | $-0.59^{***}$ | $-0.62^{***}$ |
| Participation coefficient | 0.14 | 0.05 | 0.04 | 0.03 | $-0.23$ | $-0.16$ | $-0.38^{***}$ | $-0.35^{**}$ |
| Subgraph centrality | $-0.20$ | $-0.05$ | $-0.05$ | 0.01 | 0.11 | 0.08 | $0.25^{*}$ | 0.21 |
| Integration, 80 nodes outside the core | $-0.02$ | $-0.11$ | $-0.14$ | 0.59 | $-0.51$ | $-0.64$ | $-0.61$ | $-0.56$ |
| Strength, 80 nodes outside the core | 0.17 | 0.21 | 0.13 | 0.01 | $-0.81$ | $-0.77$ | $-0.31$ | $-0.47$ |
| Integration vs strength, corrected $p$ | 0.592 | $< 0.001$ | $< 0.001$ | $< 0.001$ | 0.135 | $< 0.001$ | 0.084 | 0.956 |
| *Spearman ρ* | | | | | | | | |
| Integration | 0.19 | $-0.27^{*}$ | $-0.32^{**}$ | $0.51^{***}$ | $-0.72^{***}$ | $-0.75^{***}$ | $-0.76^{***}$ | $-0.86^{***}$ |
| Segregation | $-0.10$ | $-0.18$ | $-0.23$ | $0.24^{*}$ | $-0.35^{**}$ | $-0.34^{**}$ | $-0.13$ | $-0.22$ |
| Strength | 0.20 | $-0.11$ | $-0.19$ | 0.22 | $-0.88^{***}$ | $-0.88^{***}$ | $-0.52^{***}$ | $-0.65^{***}$ |
| Betweenness | 0.19 | 0.04 | $-0.06$ | 0.16 | $-0.64^{***}$ | $-0.66^{***}$ | $-0.39^{***}$ | $-0.51^{***}$ |
| Closeness | 0.15 | 0.03 | $-0.05$ | 0.12 | $-0.70^{***}$ | $-0.72^{***}$ | $-0.62^{***}$ | $-0.61^{***}$ |
| Participation coefficient | 0.16 | 0.06 | 0.07 | 0.02 | $-0.27^{*}$ | $-0.28^{*}$ | $-0.42^{***}$ | $-0.33^{**}$ |
| Subgraph centrality | $-0.25^{*}$ | $-0.14$ | $-0.15$ | 0.11 | 0.14 | 0.13 | $0.25^{*}$ | 0.16 |

**Table S2. Nodes with the largest effect for each perturbation.** The ten nodes with the largest effect of each perturbation, ranked from the largest. Lesion: nodes whose removal moved the CNT-fitted model furthest from the CNT data or closest to the MCS and UWS data. Stimulation: nodes whose excitation brought the UWS-fitted model closest to the CNT data (best fit over the values of $C$ with $J_{FIC} \geq 0$). Top: jump lengths, compared with the Kolmogorov–Smirnov (KS) distance; bottom: cluster occupancies, compared with the Kullback–Leibler (KL) divergence. Column headers give the group whose empirical data the model was compared with. Bold: the 10 high-integration nodes; the last row of each block gives how many of them are among the ten.

| | | Lesion (CNT-fitted model) | | Stimulation (UWS-fitted model) |
|---|---|---|---|---|
| Rank | CNT | MCS | UWS | CNT |
| *Jump lengths (KS distance)* | | | | |
| 1 | **Precuneus R** | **Cuneus L** | **Precuneus L** | Occipital Mid L |
| 2 | Frontal Med Orb R | **Cuneus R** | **Occipital Sup L** | **Occipital Sup L** |
| 3 | Frontal Sup Orb R | **Calcarine R** | **Calcarine R** | **Cingulum Post L** |
| 4 | Putamen R | **Cingulum Post R** | **Cuneus R** | **Cuneus L** |
| 5 | Rectus L | **Precuneus L** | **Precuneus R** | **Cingulum Post R** |
| 6 | Hippocampus R | Occipital Mid L | **Cuneus L** | **Calcarine L** |
| 7 | **Occipital Sup L** | **Precuneus R** | **Cingulum Post R** | **Cuneus R** |
| 8 | Occipital Inf R | Occipital Inf L | **Cingulum Post L** | **Precuneus L** |
| 9 | Frontal Sup Medial L | **Calcarine L** | Temporal Mid R | **Precuneus R** |
| 10 | Lingual R | Temporal Mid R | Frontal Inf Oper L | **Occipital Sup R** |
| *High-integration nodes* | 2/10 | 7/10 | 8/10 | 9/10 |
| *Cluster occupancies (KL divergence)* | | | | |
| 1 | **Precuneus L** | **Cingulum Post L** | **Cingulum Post R** | **Cingulum Post L** |
| 2 | **Precuneus R** | **Cingulum Post R** | **Precuneus R** | **Cingulum Post R** |
| 3 | **Calcarine R** | **Precuneus R** | **Precuneus L** | **Precuneus R** |
| 4 | **Cingulum Post R** | **Cuneus R** | **Cuneus R** | **Precuneus L** |
| 5 | **Cuneus R** | **Cuneus L** | **Cingulum Post L** | **Calcarine L** |
| 6 | **Calcarine L** | **Occipital Sup L** | **Calcarine R** | **Calcarine R** |
| 7 | **Cuneus L** | **Calcarine R** | **Cuneus L** | **Occipital Sup L** |
| 8 | **Occipital Sup L** | **Calcarine L** | **Calcarine L** | **Cuneus L** |
| 9 | **Cingulum Post L** | **Occipital Sup R** | **Occipital Sup L** | **Cuneus R** |
| 10 | **Occipital Sup R** | **Precuneus L** | **Occipital Sup R** | **Occipital Sup R** |
| *High-integration nodes* | 10/10 | 10/10 | 10/10 | 10/10 |
